# Assessing the Utility and Accuracy of ICD10-CM Nontraumatic Subarachnoid Hemorrhage Codes for Intracranial Aneurysm Research

**DOI:** 10.1101/2020.05.26.117523

**Authors:** Christopher Roark, Melissa P. Wilson, Sheila Kubes, David Mayer, Laura K. Wiley

## Abstract

**Background:** The 10th revision of International Classification of Disease, Clinical Modification (ICD10-CM) increased the number of codes to identify nontraumatic subarachnoid hemorrhage from one to twenty-two. ICD10-CM codes are able to specify the location of aneurysms causing subarachnoid hemorrhage (aSAH), however it is not clear how frequently or accurately these codes are being used in practice.

**Objective:** To systematically evaluate the usage and accuracy of location-specific ICD10-CM codes for aSAH.

**Methods:** We extracted all uses of ICD10-CM codes for nontraumatic subarachnoid hemorrhage (I60.x) during the first three years following the implementation of ICD10-CM from the billing module of the EHR for UCHealth. For those codes that specified aSAH location (I60.0-I60.6), EHR documentation was reviewed to determine whether there was an active aSAH, any patient history of aSAH, or unruptured intracranial aneurysm/s and the locations of those outcomes.

**Results:** Between October 1, 2015 – September 30, 2018, there were 3,119 instances of nontraumatic subarachnoid hemorrhage ICD10-CM codes (I60.00-I60.9), of which 297 (9.5%) code instances identified aSAH location (I60.0-I60.6). These codes accurately identified current aSAH (64%), any patient history of aSAH (84%), and any patient history of intracranial aneurysm (87%). The accuracy of identified outcome location was 53% in current aSAH, 72% for any history of aSAH, and 76% for any history of an intracranial artery.

**Conclusions:** Researchers should use ICD10-CM codes with caution when attempting to detect active aSAH and/or aneurysm location.

## INTRODUCTION

The lure of ‘big data’ for research into neurosurgical diseases is strong. Utilization of claims-based data represents a leap forward in our ability to study less prevalent diseases like aneurysmal subarachnoid hemorrhage (aSAH). Although roughly 6.5-9.7 million people in the United States harbor an unruptured intracranial aneurysm, only about 30,000 of those aneurysms rupture in a given year.^1–3^ The rupture of an intracranial aneurysm causes an aSAH which has a high (50%) mortality rate and often leaves survivors with permanent disability.^4^ Conducting traditional randomized controlled trials (RCTs) are challenging for aSAH due to the low prevalence of the condition. For example, the International Subarachnoid Aneurysm Trial (ISAT), one of the largest randomized controlled trials in neurosurgery, required seven years to recruit a cohort of 2,143 patients across 43 centers in Europe and North America.^5^ While RCTs are the gold standard for clinical evidence, the large sample size of administrative claims data is alluring for real-world evidence generation.

According to a systematic review of claims-based research in neurosurgery, 50 studies of aneurysms and aSAH had been published in the top three neurosurgery journals between 2000 and 2016. One of the major limitations identified across these studies was the lack of specific clinical data required for appropriate risk adjustment.^6^ Claims databases often use International Classification of Disease, 9th Revision, Clinical Modification (ICD9-CM) codes to identify clinical diagnosis. Unfortunately, ICD9-CM only has a single code for non-traumatic subarachnoid hemorrhage (430), which includes both aSAH and spontaneous hemorrhages that do not harbor aneurysms. Importantly, aneurysm location (which is an important risk factor for aneurysm rupture and correlates with aSAH outcome^7–10^), is not captured by this code. These data are not typically available in large claims databases, reducing the impact and value of these databases for aSAH research.

However, on October 1, 2015, the United States began using the 10th revision of ICD codes (ICD10-CM). This new claims list dramatically expanded the number of codes for nontraumatic subarachnoid hemorrhage from one to twenty-two. These new codes give a specific location for common sites of ruptured intracranial aneurysms, which represents a significant improvement in the utility of administrative claims databases for aSAH research. However, the accuracy of these codes to identify aSAH or aSAH location has not yet been determined.

In this study, we sought to evaluate the overall usage and accuracy of location-specific ICD10-CM codes for aSAH. We examined all billing uses of ICD10-CM codes for non-traumatic hemorrhage in the first three years post-implementation at a large multi-hospital healthcare system located across the front range of Colorado. Detailed chart review was performed to verify aSAH status and aneurysm rupture location. These data should provide insight into the utility of large ICD10-CM based claims databases for aSAH research.

## METHODS

### Study Cohort

We extracted all uses of ICD10-CM codes for nontraumatic subarachnoid hemorrhage (I60.x) during the first three years following the implementation of ICD10-CM (October 1, 2015 – September 30, 2018) from the billing module of the electronic health record (EHR) for UCHealth, a healthcare system in Colorado consisting of 13 hospitals and multiple outpatient clinics. This study only included codes used for patients 18 years of age or older.

### Assess utility of codes for intracranial aneurysm research

We calculated the frequency of each ICD10-CM code used for billing (e.g., every “code instance”), “unique visit” (code/s billed on the same day for the same patient), and “unique patient” (code/s ever billed for the same patient).

### Assess accuracy of codes for intracranial aneurysm research

To determine the accuracy of the code used, CR, SK and/or MW reviewed progress notes, imaging, procedure reports and patient history from the EHR for each visit billed with an ICD10-CM code that specified an artery location (I60.0-I60.6). Reviewers determined whether there was active management of an aSAH at that code instance date. “Active aSAH” was defined as the hospitalization when an aneurysm rupture was diagnosed/treated, an immediate hospitalization or inpatient rehabilitation following the diagnosis/treatment (at the same or different facility), or acute follow-up visits within 6 weeks post-discharge. Reviewers then also considered the patient’s entire chart to determine whether they had ever experienced an aSAH (“History of aSAH”) or were diagnosed with an aneurysm (“History of Aneurysm”). The location/s of all identified active aSAH and historical aSAH/aneurysms were also recorded.

Using the results from the chart review, we first calculated the percent accuracy of each code instance identifying an active aSAH, history of aSAH, or history of aneurysm outcome regardless of specified location. Any code instance that had no occurrence of the specified outcome was classified as a “false positive”. For each outcome (aSAH, history of aSAH, and history of aneurysm), we then assessed the accuracy of aneurysm location identified by the ICD10-CM code instance. Code instances where an aneurysm location could not be identified from the medical record were removed from this analysis. For codes that specified an artery but where laterality was unspecified (i.e., I60.00, I60.10, I60.30, I60.50), outcomes that occurred in that artery were considered “correct”. For codes that specified the laterality of an artery (e.g., I60.01, I60.02, I60.11, I60.12, I60.31, I60.32, I60.51, I60.52), outcomes that occurred in the artery but at the alternative laterality were classified as incorrect (wrong side). Code instances for I60.6 - “Other Artery” were classified as incorrect (wrong artery) if there was an available ICD10-CM code for the actual artery location of the outcome (e.g., carotid siphon & bifurcation, middle cerebral artery, anterior communicating artery, posterior communicating artery, basilar artery, or vertebral artery). For patients with multiple aneurysms, accuracy was assigned manually for each outcome. Active aSAH and history of aSAH outcomes were only considered correct for the ruptured artery/bleeding source. For patients with multiple historical sites of aSAH and/or aneurysms, we assigned status conservatively giving the most credit possible (i.e., accurate > incorrect - wrong side > incorrect - wrong artery).

We also analyzed whether ICD10-CM code occurrences could be used to identify patients with any history of multiple aneurysms (ruptured or unruptured). First we tested whether patients who had two or more distinct ICD10-CM codes across their record had a history of multiple aneurysms. Then we considered the more specific condition: whether patients who explicitly had two or more arteries identified (i.e., two or more I60.0-I60.6 codes) had a history of multiple aneurysms.

Chi-square or Fisher’s exact tests were performed to evaluate for group differences, using a significance level of p<0.05. No adjustment was made for multiple testing, as outcomes were considered complementary. We also calculated the positive predictive value for both definitions.

### Characterize usage of codes over time

To understand whether ICD10-CM code usage changed over time, we compared the frequency of codes that specified an artery location (I60.0-I60.6) to those that did not (I60.7-I60.9) each month over the study period. Similarly, we compared the accuracy of codes that specified an artery location (I60.0-I60.6) monthly over the study period.

Study data were collected and managed using REDCap electronic data capture tools hosted at University of Colorado Anschutz Medical Campus.^11^ All analyses were performed using R version 3.6.0^12^ and a variety of packages for data processing, graphing, and reporting.^13–17^ This study was reviewed and approved by the Colorado Multiple Institutional Review Board (COMIRB).

## RESULTS

Between October 1, 2015 – September 30, 2018, there were 3,119 instances of nontraumatic subarachnoid hemorrhage ICD10-CM codes (I60.00-I60.9) billed in Epic across UCHealth. These codes represented 1,408 unique patients and 2,902 unique visits.

The majority of these code instances (2,822, 90.5%) did not specify an aSAH location (i.e., I60.7-I60.9). The 297 (9.5%) codes that identified aSAH location (I60.0-I60.6) represented 199 unique patients and 279 unique visits. A complete summary of code instances and unique patient frequencies by code is available in **Figure 1.** The highest frequency code was I60.2 (Nontraumatic subarachnoid hemorrhage from anterior communicating artery), and across all location-specific codes where laterality was identified, right lateralities accounted for 65% of cases. Of the ICD10-CM codes that had designations for artery laterality available (i.e., I60.0*, I60.1*, I60.3*, I60.5*) 10.7% of code instances did not specify laterality. **Table 1** contains usage frequencies for all nontraumatic SAH codes over the study period.

**Table 1.**
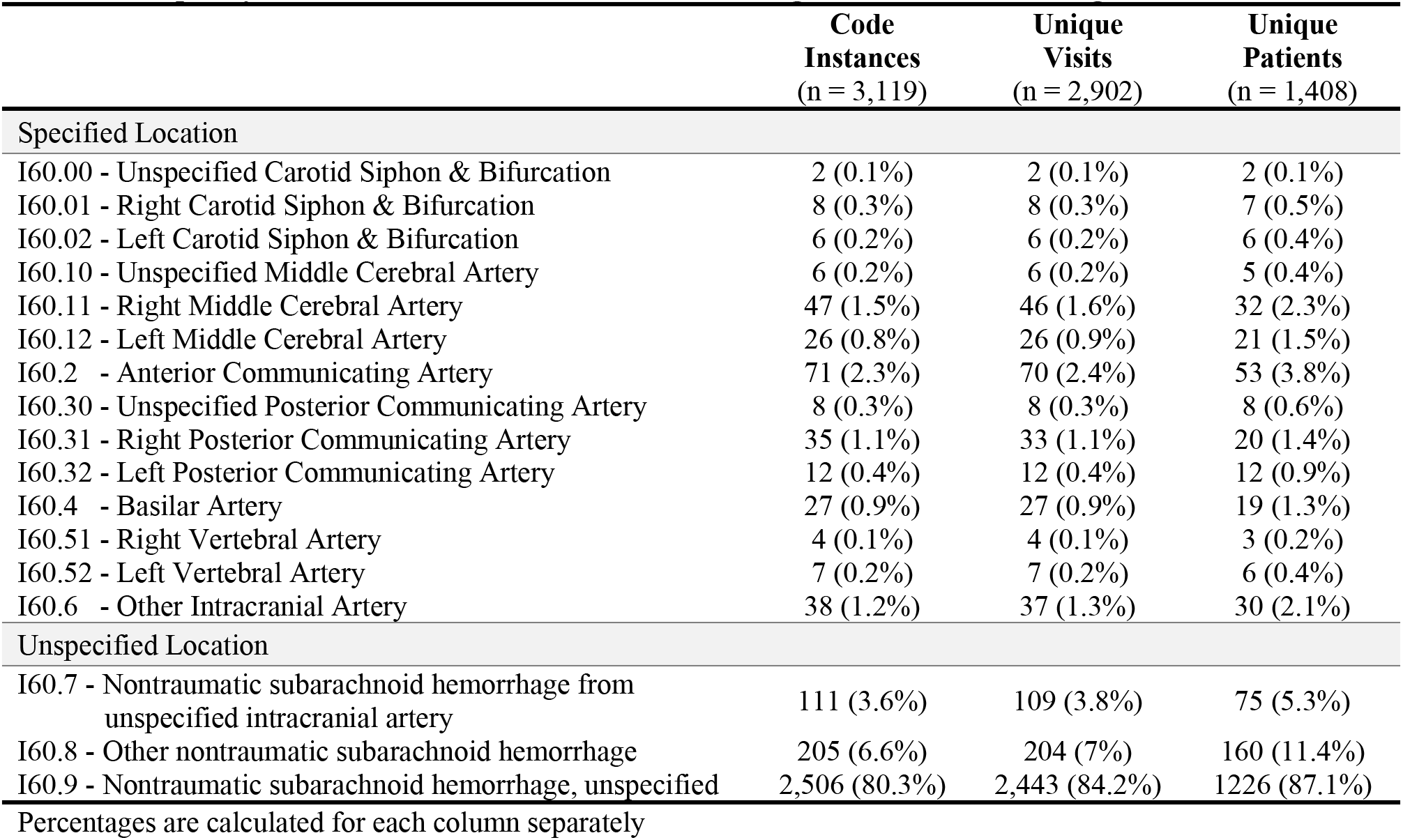
Frequency of Nontraumatic Subarachnoid Hemorrhage ICD10-CM Code Usage from 10/2015-9/2018.

|  | **Code**<br>**Instances**<br>**(n = 3,119)** | **Unique**<br>**Visits**<br>**(n = 2,902)** | **Unique**<br>**Patients**<br>**(n = 1,408)** |
| --- | --- | --- | --- |
| **Specified Location** |  |  |  |
| I60.00 - Unspecified Carotid Siphon & Bifurcation | 2 (0.1%) | 2 (0.1%) | 2 (0.1%) |
| I60.01 - Right Carotid Siphon & Bifurcation | 8 (0.3%) | 8 (0.3%) | 7 (0.5%) |
| I60.02 - Left Carotid Siphon & Bifurcation | 6 (0.2%) | 6 (0.2%) | 6 (0.4%) |
| I60.10 - Unspecified Middle Cerebral Artery | 6 (0.2%) | 6 (0.2%) | 5 (0.4%) |
| I60.11 - Right Middle Cerebral Artery | 47 (1.5%) | 46 (1.6%) | 32 (2.3%) |
| I60.12 - Left Middle Cerebral Artery | 26 (0.8%) | 26 (0.9%) | 21 (1.5%) |
| I60.2 - Anterior Communicating Artery | 71 (2.3%) | 70 (2.4%) | 53 (3.8%) |
| I60.30 - Unspecified Posterior Communicating Artery | 8 (0.3%) | 8 (0.3%) | 8 (0.6%) |
| I60.31 - Right Posterior Communicating Artery | 35 (1.1%) | 33 (1.1%) | 20 (1.4%) |
| I60.32 - Left Posterior Communicating Artery | 12 (0.4%) | 12 (0.4%) | 12 (0.9%) |
| I60.4 - Basilar Artery | 27 (0.9%) | 27 (0.9%) | 19 (1.3%) |
| I60.51 - Right Vertebral Artery | 4 (0.1%) | 4 (0.1%) | 3 (0.2%) |
| I60.52 - Left Vertebral Artery | 7 (0.2%) | 7 (0.2%) | 6 (0.4%) |
| I60.6 - Other Intracranial Artery | 38 (1.2%) | 37 (1.3%) | 30 (2.1%) |
| **Unspecified Location** |  |  |  |
| I60.7 - Nontraumatic subarachnoid hemorrhage from unspecified intracranial artery | 111 (3.6%) | 109 (3.8%) | 75 (5.3%) |
| I60.8 - Other nontraumatic subarachnoid hemorrhage | 205 (6.6%) | 204 (7%) | 160 (11.4%) |
| I60.9 - Nontraumatic subarachnoid hemorrhage, unspecified | 2,506 (80.3%) | 2,443 (84.2%) | 1226 (87.1%) |

Percentages are calculated for each column separately

**Figure 1.**
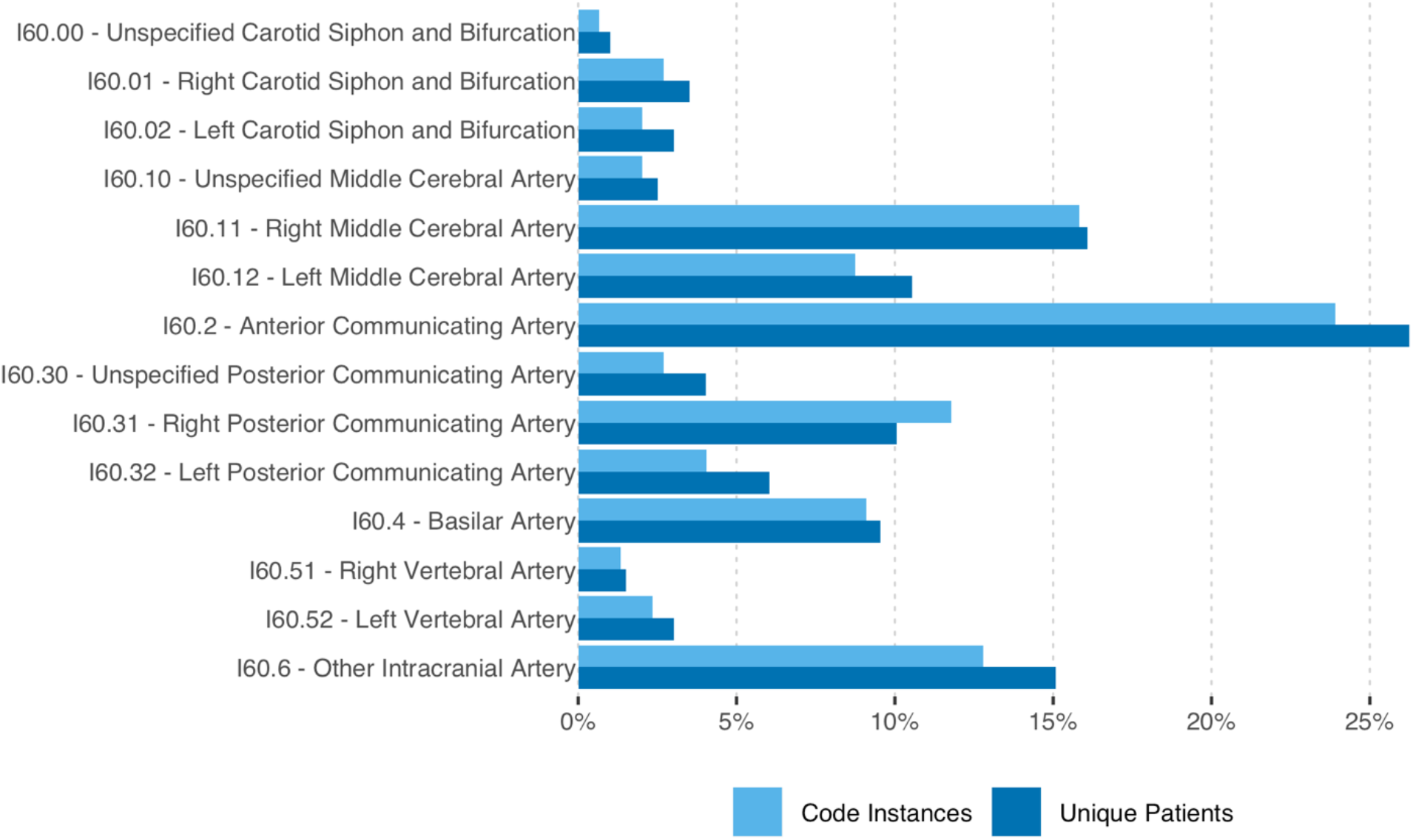
Frequency of ICD10-CM codes that specify location of nontraumatic subarachnoid hemorrhage. The usage of each ICD10-CM code that specifies aSAH location (i.e., I60.0-I60.6) is provided as a percent of the total code instances (light blue) or unique patients (dark blue).

Chart review revealed that out of all location-specific code instances, no evidence of an aSAH (current or historical) could be found in 48 (16.3%), and 35 (11.9%) had no evidence of any intracranial aneurysm. In the cases where no aneurysm was documented, 18 had a current non-aneurysmal SAH (non-traumatic), and 17 had neither an SAH (of any origin) nor an aneurysm. In two code instances, a distant history of multiple aSAHs was documented, but location could not be determined; these instances were dropped from the remaining analyses. A summary of the accuracy rate for all three tested outcomes (current aSAH, history of aSAH, and history of aneurysm) is presented in **Figure 2**.

**Figure 2.**
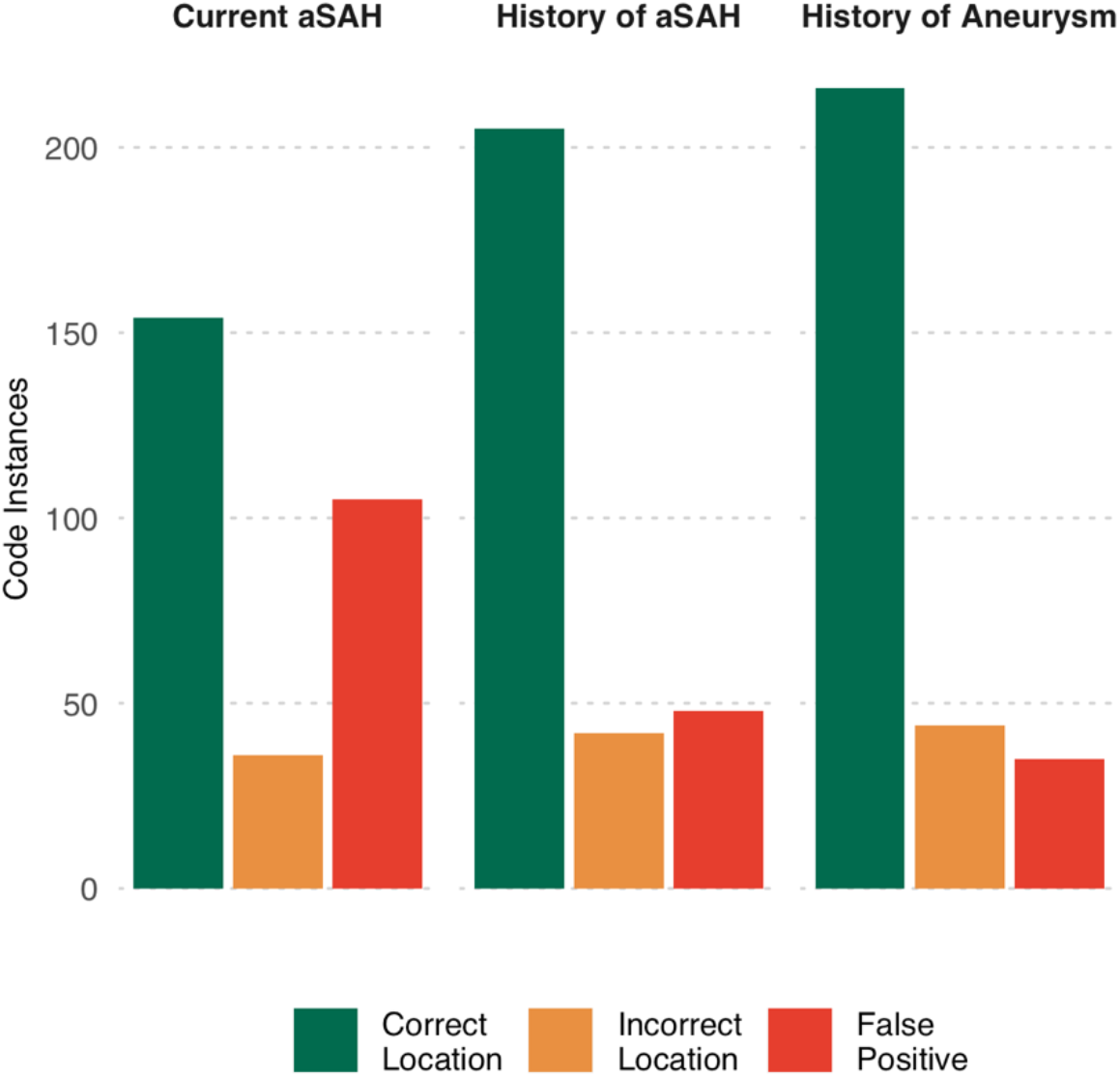
Accuracy of ICD10-CM that specify location of nontraumatic subarachnoid hemorrhage to identify current aneurysmal subarachnoid hemorrhage, any history of subarachnoid hemorrhage, or any history of intracranial aneurysm. The total number of code instances that correctly identified both the outcome (current aSAH - left, history of aSAH - middle, history of intracranial aneurysm - right) and artery is displayed in green. The total number of code instances that correctly identified the outcome but not the correct artery is displayed in orange. The total number of code instances that did not correctly identify the outcome are displayed in red.

### Ability to Detect Current Aneurysmal Subarachnoid Hemorrhage

Of the codes that specified artery location, 190 (64.4%) were entered during an “active/current” aSAH episode. Over half of the code instances identified the correct aSAH location with 154 (52.2%) with an exact match and 4 (1.4%) that indicated the correct artery location but did not specify laterality. Of the incorrect instances, 1 (0.3%) identified the correct artery but incorrect laterality, while 31 (10.5%) identified the wrong artery. Accuracy of individual ICD10-CM codes to detect active aSAH are available in **Table 2.**

**Table 2.** Accuracy to Detect Current Aneurysmal Subarachnoid Hemorrhage.

|  | Correct | Incorrect |  | False Positive |  |
| --- | --- | --- | --- | --- | --- |
|  |  | Wrong<br>Laterality | Wrong<br>Artery | Non-aneurysmal<br>SAH | No SAH |
| I60.00 - Unspecified Carotid Siphon & Bifurcation | - | - | - | - | 2 (100%) |
| I60.01 - Right Carotid Siphon & Bifurcation | 6 (75%) | - | - | - | 2 (25%) |
| I60.02 - Left Carotid Siphon & Bifurcation | 1 (16.7%) | - | 2 (33.3%) | - | 3 (50%) |
| I60.10 - Unspecified Middle Cerebral Artery | 2 (33.3%) | - | - | - | 4 (66.7%) |
| I60.11 - Right Middle Cerebral Artery | 18 (38.3%) | - | 3 (6.4%) | 4 (8.5%) | 22 (46.8%) |
| I60.12 - Left Middle Cerebral Artery | 13 (50%) | - | - | 6 (23.1%) | 7 (26.9%) |
| I60.2 - Anterior Communicating Artery | 47 (66.2%) | - | 7 (9.9%) | - | 17 (23.9%) |
| I60.30 - Unspecified Posterior Communicating Artery | 2 (25%) | - | 1 (12.5%) | - | 5 (62.5%) |
| I60.31 - Right Posterior Communicating Artery | 21 (60%) | - | 3 (8.6%) | - | 11 (31.4%) |
| I60.32 - Left Posterior Communicating Artery | 8 (66.7%) | 1 (8.3%) | 3 (25%) | - | - |
| I60.4 - Basilar Artery | 16 (59.3%) | - | 2 (7.4%) | 3 (11%) | 6 (22.2%) |
| I60.51 - Right Vertebral Artery | 3 (75%) | - | 1 (25%) | - | - |
| I60.52 - Left Vertebral Artery | 4 (57.1%) | - | 2 (28.6%) | - | 1 (14.3%) |
| I60.6 - Other Intracranial Artery | 17 (47.2%) | - | 7 (19.4%) | 5 (13.9%) | 7 (19.4%) |

Percentages are calculated for each code separately.

### Ability to Detect Any History of Aneurysmal Subarachnoid Hemorrhage

Of the codes that specified artery location, 247 (83.7%) were used for patients who had an aSAH at any point in their history. The majority (n = 205, 69.5%) correctly identified the location of the current or previous aSAH with 7 (2.4%) indicating the correct artery location without specifying artery laterality. Of those that indicated the incorrect location, 1 (0.3%) identified the correct artery, but indicated the wrong side, and 34 (11.5%) had the wrong artery. In 13 code instances (4.4%), the patient had one or more aneurysms, but no history of an aSAH. Accuracy of individual ICD10-CM codes to detect any history of aSAH are available in **Table 3.**

**Table 3.** Accuracy to Detect Current Any History of Aneurysmal Subarachnoid Hemorrhage.

|  | Correct | Incorrect |  | False Positive |
| --- | --- | --- | --- | --- |
|  |  | Wrong<br>Laterality | Wrong<br>Artery | No History of<br>aSAH |
| I60.00 - Unspecified Carotid Siphon & Bifurcation | - | - | - | 2 (100%) |
| I60.01 - Right Carotid Siphon & Bifurcation | 7 (87.5%) | - | - | 1 (12.5%) |
| I60.02 - Left Carotid Siphon & Bifurcation | 2 (33.3%) | - | 2 (33.3%) | 2 (33.3%) |
| I60.10 - Unspecified Middle Cerebral Artery | 3 (50%) | - | - | 3 (50%) |
| I60.11 - Right Middle Cerebral Artery | 30 (63.8%) | - | 3 (6.4%) | 14 (29.8%) |
| I60.12 - Left Middle Cerebral Artery | 18 (69.2%) | - | - | 8 (30.8%) |
| I60.2 - Anterior Communicating Artery | 61 (85.9%) | - | 8 (11.3%) | 2 (2.8%) |
| I60.30 - Unspecified Posterior Communicating Artery | 4 (50%) | - | 2 (25%) | 2 (25%) |
| I60.31 - Right Posterior Communicating Artery | 31 (88.6%) | - | 3 (8.6%) | 1 (2.8%) |
| I60.32 - Left Posterior Communicating Artery | 8 (66.7%) | 1 (8.3%) | 3 (25%) | - |
| I60.4 - Basilar Artery | 20 (74.1%) | - | 3 (11.1%) | 4 (14.8%) |
| I60.51 - Right Vertebral Artery | 3 (75%) | - | 1 (25%) | - |
| I60.52 - Left Vertebral Artery | 4 (57.1%) | - | 2 (28.6%) | 1 (14.3%) |
| I60.6 - Other Intracranial Artery | 21 (58.3%) | - | 7 (19.4%) | 8 (22.2%) |

Percentages are calculated for each code separately.

### Ability to Detect Any History of Intracranial Aneurysm

The majority of location-specific aSAH code instances (n = 260, 88.1%) accurately identified patients who had any history of an intracranial aneurysm at the time of entry. The location accuracy rate was highest among the three outcomes with 216 (73.2%) having an exact location match and an additional 9 (3%) that identified the correct artery but did not specify that aneurysm laterality. The majority of the incorrect locations indicated the wrong artery (n = 34, 11.5%), while 1 (0.3%) identified the correct artery, but indicated the wrong laterality. Accuracy of individual ICD10-CM codes to detect any history of one or more intracranial aneurysm are available in **Table 4.**

**Table 4.** Accuracy to Detect Current Any History of Aneurysm.

|  | Correct | Incorrect |  | False Positive |
| --- | --- | --- | --- | --- |
|  |  | Wrong Laterality | Wrong Artery | No History of Aneurysm |
| I60.00 - Unspecified Carotid Siphon & Bifurcation | - | - | - | 2 (100%) |
| I60.01 - Right Carotid Siphon & Bifurcation | 8 (100%) | - | - | - |
| I60.02 - Left Carotid Siphon & Bifurcation | 2 (33.3%) | - | 2 (33.3%) | 2 (33.3%) |
| I60.10 - Unspecified Middle Cerebral Artery | 5 (83.3%) | - | - | 1 (16.7%) |
| I60.11 - Right Middle Cerebral Artery | 34 (72.3%) | 1 (2.1%) | 3 (6.4%) | 9 (19.1%) |
| I60.12 - Left Middle Cerebral Artery | 18 (69.2%) | - | - | 8 (30.8%) |
| I60.2 - Anterior Communicating Artery | 65 (91.5%) | - | 6 (8.5%) | - |
| I60.30 - Unspecified Posterior Communicating Artery | 4 (50%) | - | 2 (25%) | 2 (25%) |
| I60.31 - Right Posterior Communicating Artery | 31 (88.6%) | - | 3 (8.6%) | 1 (2.8%) |
| I60.32 - Left Posterior Communicating Artery | 9 (75%) | - | 3 (25%) | - |
| I60.4 - Basilar Artery | 20 (74.1%) | - | 3 (11.1%) | 4 (14.8%) |
| I60.51 - Right Vertebral Artery | 3 (75%) | - | 1 (25%) | - |
| I60.52 - Left Vertebral Artery | 4 (57.1%) | - | 2 (28.6%) | 1 (14.3%) |
| I60.6 - Other Intracranial Artery | 22 (61.1%) | - | 9 (25%) | 5 (13.9%) |

Percentages are calculated for each code separately.

### Ability to Detect Multiple Aneurysms

Of those individuals with a location specified aSAH code, 35 (18%) had a history of multiple aneurysms, representing 76 total code instances. The association between having two or more distinct SAH ICD10-CM codes across their record and chart review revealing a history of multiple aneurysm was not significant (Ꭓ^2^ = 1.12, p = 0.29) and had a positive predictive value of 20.3%. Comparing the more strict criteria of having two or more arteries identified (i.e., two of more I60.0-I60.6 codes) associated with a history of multiple aneurysms was also not significant (Fisher’s Exact p-value = 0.08, positive predictive value = 31.6%).

### Coding Usage and Accuracy Over Time

During the first year after implementation of ICD10-CM, 47 location-specific aSAH codes were entered; 111 and 139 codes were entered the second and third years, respectively. The majority of codes across all three years did not specify location (94.3% year 1, 89.5% year 2, 88.8% year 3). **Figure 3** shows the frequency of monthly code use over the study period. Among those code instances that specified an aneurysm location the accuracy of the location specified was 65.2% in the first year and improved to 72.3% and 76.6% in the second and third years. **Figure 4** shows the monthly use and accuracy of location-specific ICD10-CM codes during the study period.

**Figure 3.**
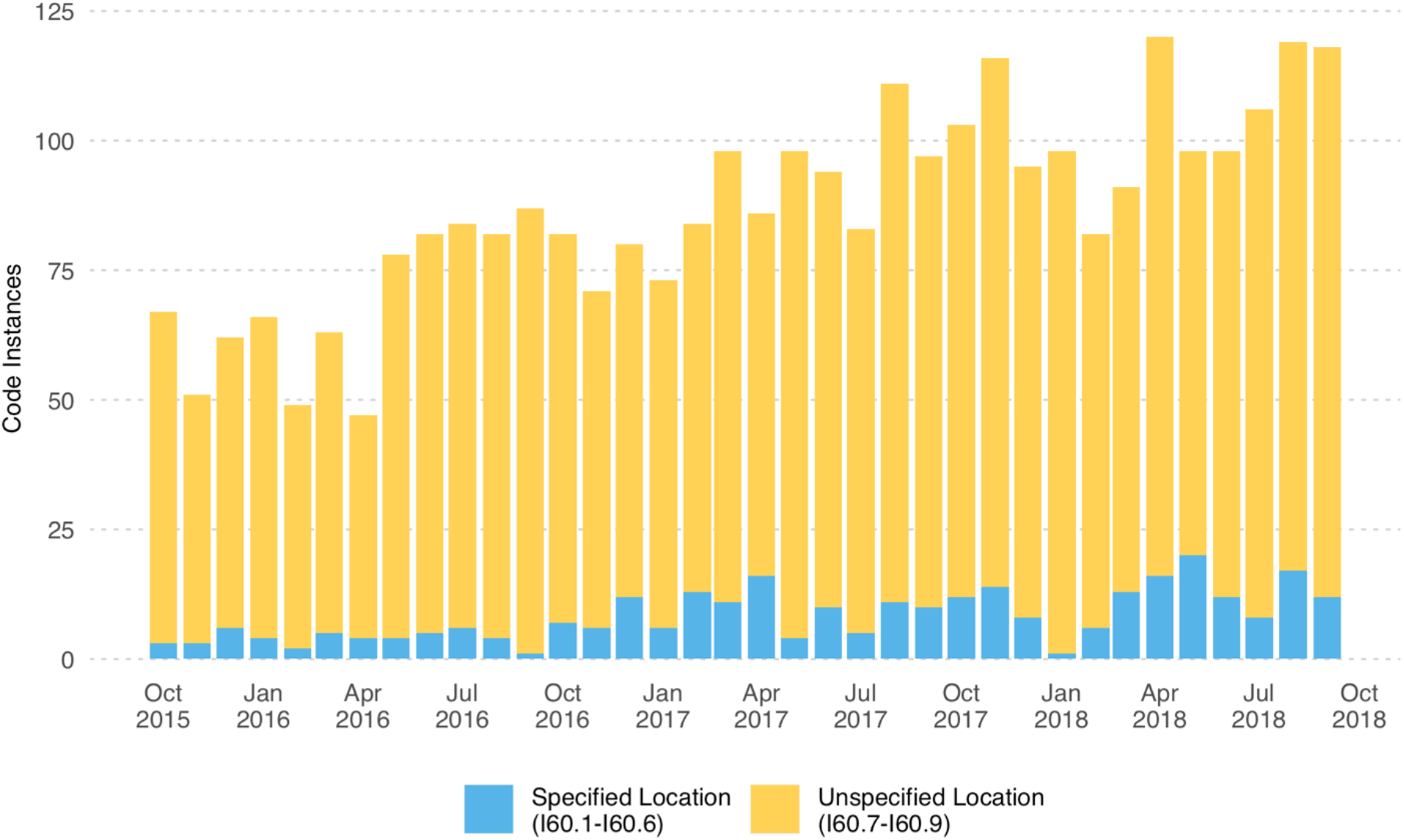
Usage frequency of ICD10-CM codes for nontraumatic subarachnoid hemorrhage across October 1, 2015 - September 30, 2018. The monthly total number of ICD10-CM code instances for nontraumatic subarachnoid hemorrhage (I60.0-I60.9) across the study period. The count of codes that specified aSAH location (I60.0-I60.6) are shown in light blue, while those that did not specify location (I60.7-I60.9) are shown in yellow.

**Figure 4.**
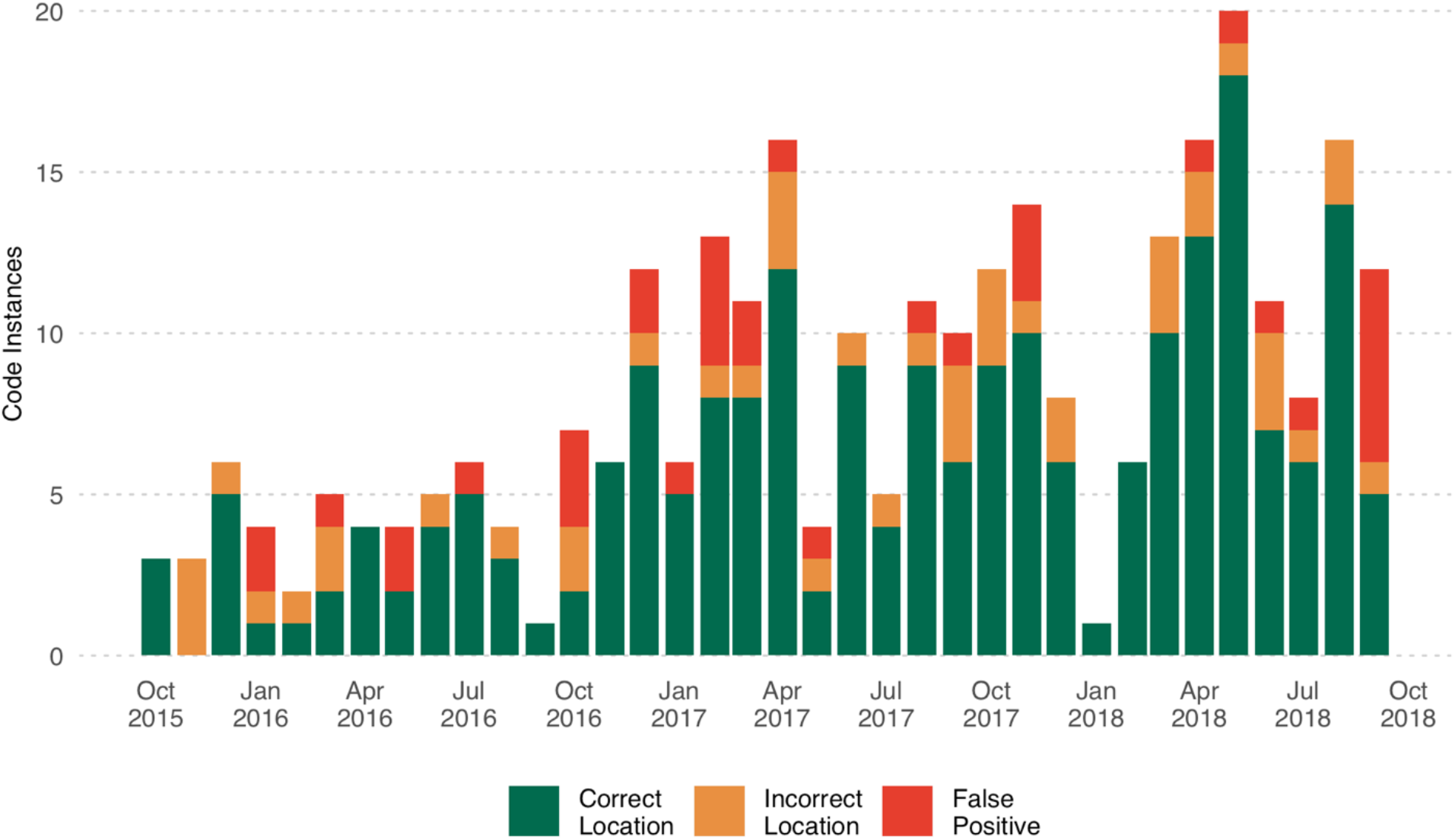
Accuracy of ICD10-CM codes that specify location of nontraumatic subarachnoid hemorrhage to detect any history of intracranial aneurysm across October 1, 2015 - September 30, 2018. The monthly total number of ICD10-CM code instances that specify location of nontraumatic subarachnoid hemorrhage (I60.0-I60.6) across the study period. The total number of code instances that correctly identified both the history of an intracranial aneurysm (ruptured or unruptured) and the artery location is displayed in green. The total number of code instances that correctly identified a history of intracranial aneurysm but not the correct artery is displayed in orange. The total number of code instances that did not correctly identify a history of intracranial aneurysm are displayed in red.

## DISCUSSION

The implementation of location-specific nontraumatic subarachnoid hemorrhage ICD10-CM codes in October of 2015 has the potential to significantly improve big data research of aSAH in the United States. Until this change, researchers using large claims databases had to trade larger study sizes for the lack of availability of important risk factors. Because aneurysm location plays such a key role in risk modeling and clinical outcome studies of intracranial aneurysm, this represented a significant limitation for their studies. While there is great promise in these new codes, there has been limited data available on how they are being used in practice and whether they accurately identify aSAH and/or aSAH location. This is the first comprehensive evaluation of the accuracy of ICD10-CM codes for nontraumatic subarachnoid hemorrhage performed in the United States.

Over the first three years of use of ICD10-CM codes at UCHealth, we found that the vast majority (90.5%) of code instances did not take advantage of the ICD10-CM codes that specified aSAH location. While the frequency of use did increase over the study period, the highest usage rate was only 11.2% in the third year of the study. If the usage patterns at our institution are representative of the broader healthcare industry, this dramatically reduces the utility of location-specific nontraumatic subarachnoid hemorrhage ICD10-CM codes for research.

When location-specific nontraumatic subarachnoid hemorrhage ICD10-CM codes *were* used, the relative frequency of aneurysm location matches what is reported in the neurosurgical literature. Over 80% of intracranial aneurysms tend to occur in the carotid circulation with the anterior communicating artery representing the most common location, followed by the posterior communicating and middle cerebral arteries. Upon review 18% of patients in our study had multiple aneurysms, which is consistent with previously reported literature.^18,19^ These data suggest that there are not obvious differences between the patient population seen at UCHealth and those represented in the scientific literature.

Although we specifically examined billing uses of the ICD10-CM codes (i.e., the diagnosis used to justify payment for healthcare services rendered), only 64% were used during an active aSAH episode. This performance is lower than international studies that validated ICD10 SAH detection rates across three hospitals in Calgary, Canada (91% correct; CI: 76-98%).^20^ This difference may be due to the different healthcare reimbursement landscapes in Canada (single payer) vs the United States. Based on our data, researchers using US claims data should be wary of interpreting ICD10-CM nontraumatic subarachnoid hemorrhage code use as indicative of an active aSAH. However, the accuracy of these codes to detect any patient history of an intracranial aneurysm was high (88.1%) suggesting that cautious use of these codes for aneurysm research may be appropriate.

When detecting any patient history of an aneurysm, the location accuracy was moderate (73.2%) and increased to 76.6% in the third year of the study. The majority of errors indicated the wrong artery location (14.9%), and these errors were not evenly distributed across arteries. Among the three most common arteries identified (middle cerebral, anterior communicating, and posterior communicating) the wrong artery was identified only 3.8% of middle cerebral artery codes, but 8.5% and 14.5% of the anterior and posterior communicating arteries respectively. Importantly, the abbreviations for these two locations (ACoA and PCoA) could easily be confused with the anterior cerebral artery (ACA), posterior inferior cerebellar artery (PICA), or posterior cerebral artery (PCA). Indeed, of the errors for these two locations, 66% of the incorrect ACoA codes were identified in chart review to be an aneurysm in the anterior cerebral artery and 37.5% of the incorrect PCoA were an aneurysm in the posterior inferior cerebellar artery. As with other hospitals, we utilize professional coders to bill inpatient encounters. Medical coders come from a variety of educational backgrounds and while a number of certifications are available, the level of domain-specific training they receive is unclear. The number of very similar abbreviations used across neurosurgical notes make these types of errors more likely and should be considered by researchers when using codes for these sites.

Although the focus of this project was on determining the accuracy of ICD10-CM codes for identifying aSAH occurrence and aSAH location, we also investigated whether these codes could be used to detect patients who harbor multiple aneurysms. Patients who have multiple aneurysms are at higher risk for aSAH.^7–10^ Unfortunately neither definition of multiple aneurysms used had sufficient resolution (PPV < 35%) for use in research.

This study is limited by the fact that it represents a single healthcare system, with the majority of reviewed records coming from a single hospital. Additionally, we only examined the first three years of data after implementation of ICD10-CM. In our data, both the usage and accuracy of these codes increased over the study period. It is not clear whether this trend will continue linearly or eventually plateau at a stable usage rate. More studies utilizing other centers as well as longitudinal studies to assess for increasing trends of specific and correct usage will help clinicians and data scientists better assess the utility of ICD10-CM codes for long-term intracranial aneurysm research.

## CONCLUSION

It is logical to assume that the newly available level of specificity of ICD10-CM will lead many researchers to incorporate aneurysm location into future clinical data science studies. Aneurysm location is an important predictor of rupture for unruptured aneurysms ^21–23^, and the location is often a surrogate for the difficulty in repair. However, our data suggest that these codes are used infrequently and have variable accuracy to detect intracranial aneurysm outcomes and location. Therefore, depending on their specific research question, researchers should use these data with caution when attempting to detect active aSAH and/or aneurysm location.

## Acknowledgements

We would like to thank Rachel Zucker for providing project management support for this project. This project was also supported by the Health Data Compass Data Warehouse project (healthdatacompass.org) and the NIH/NCRR Colorado CTSI Grant Number UL1 RR02578. It’s contents are the authors’ sole responsibility and do not necessarily represent official NIH views.

